# Supercoiling twists Cas9 off-target discrimination when nicking and cleaving

**DOI:** 10.64898/2026.04.01.715559

**Authors:** Ieva Jaskovikaitė, Hidde S. Offerhaus, Marius Vinogradovas, Uršulė Barkauskaitė, Martin Depken, Stephen K. Jones

## Abstract

Programmed with an RNA guide, Cas9 nuclease directs double-strand DNA cleavage via its two nuclease domains. However, Cas9 sometimes falsely identifies DNA targets by binding and cleaving DNA that does not match its PAM and guide RNA sequence requirements. Cas9’s specificity is often affected by DNA topology, as DNA negative supercoiling can increase off-target activity while positive supercoiling can even prevent on-target activity. Such dramatic DNA topological changes routinely occur in cells as a result of transcription and replication, making Cas9’s specificity a challenge for gene editing. To determine how Cas9 imparts its specificity across sequences and topologies, we directly mapped kinetic cleavage rates and sites for thousands of relaxed and negatively-supercoiled target sequences via NucleaSeq. We find that: Negative supercoiling can accelerate off-target cleavage a thousand-fold, and shift cleavage sites by two nucleotides. Guide-target mispairs differently impact RuvC and HNH domains, which can lead to topology-dependent nicking by Cas9. Finally, we predict these variations in Cas9 cleavage activity with a biophysical model that accounts for DNA topological state. These efforts expose Cas9’s strand-specific off-target cleavage activity and can improve off-target identification for more predictable and safer gene editing.

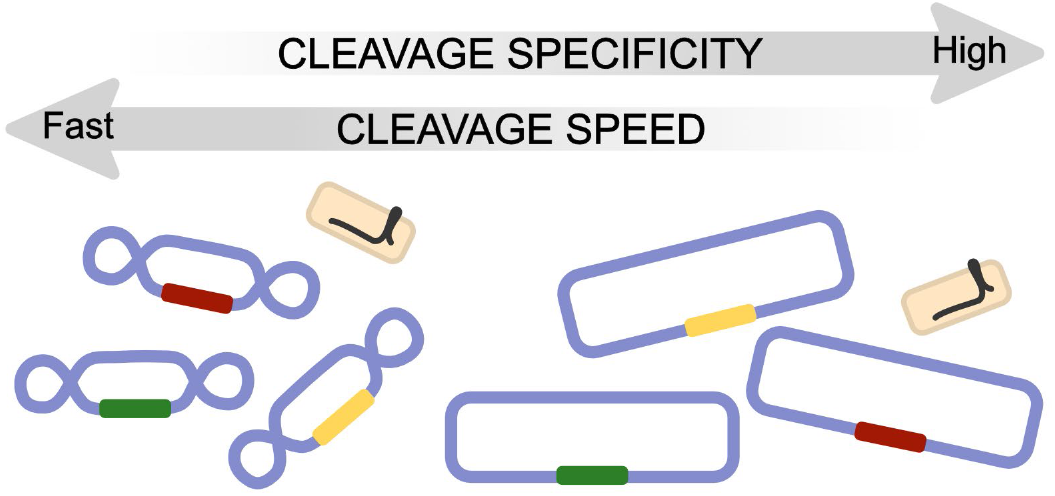

## Introduction

The *Streptococcus pyogenes* Clustered Regularly Interspaced Short Palindromic Repeats (CRISPR) – associated 9 (Cas9) nuclease forms a ribonucleoprotein (RNP) complex with a guide RNA (gRNA; made of CRISPR-RNA and *trans*-activating RNA)^1^, that is capable of recognizing target DNAs^2^. Upon locating an NGG protospacer adjacent motif (PAM), Cas9 forms an R-loop by melting the adjacent DNA and pairing it with its gRNA^2–4^. Given sufficient complementarity, the two nuclease domains, HNH and RuvC, cleave target and non-target DNA strands, respectively^5,6^. Due to simple gRNA reprogramming of Cas9 to different targets, the nuclease is widely used in the lab and attractive for the clinic^7–9^.

Cas9 occasionally operates on partially-matched targets, which can trigger edits at unintended locations and even chromosomal rearrangements^10,11^. Such lack of specificity includes accepting non-canonical PAM sequences (*i*.*e*. NGA and NAG)^12,13^ and mispairs within the 20-nucleotide target^14–20^. Cas9 tolerates gRNA-relative mispairs *in vitro* and *in vivo*^14–20^, with greater tolerance for PAM-distal mispairs than for PAM-proximal ‘seed’ mispairs (positions ∼5-10) ^14^. Cas9’s cumulative activity across off-targets can be as frequent as it is for on-targets^17,19,21^. For gRNAs tested on the human genome, single base mismatches are most prevalent within off-targets^21^, even within off-targets containing other mispairs (gRNA-relative deletions or insertions)^19^. Concerns about off-targets have motivated investigations into off-target binding patterns^15,22^, cleavage patterns^15,17^, and their relation to gRNA sequence^23–25^. These studies, and the models built upon them^26–32^, help make genome editing safer by pinpointing, predicting and avoiding Cas9’s promiscuity. Unfortunately, current *in silico* prediction models still miss *bona fide* off-targets, and would benefit from including additional variables that impact Cas9 activity.

Recent studies show that DNA topology – specifically negative supercoiling (nSC) – can increase Cas9 off-target cleavage^17,33–35^, and processes like transcription induce such topological changes^36^. Negative supercoiling induces transient DNA unwinding^37,38^ and increases Cas9’s propensity for cleaving genomic off-targets^17,35^. Here, negative torque on the DNA stabilizes R-loop intermediates for genomic off-targets with PAM-distal mispairs, encouraging cleavage. However, PAM-proximal mispairs discourage R-loop intermediate formation, and negative torque provides insufficient assistance to overcome them^34^. Similarly, catalytically “dead” Cas9 (dCas9) can dwell longer on off-targets that are underwound^33,34^. Yet, we still miss a clear understanding of the role that supercoiling plays across Cas9 binding and cleavage in the wider off-target space (in relation to mispair types, identities and positions), rather than mismatch mispairs alone^39^. We also understand that Cas9 cuts relaxed targets at different positions (and even trims ends) depending on target sequence^6,15,40^. However, we similarly miss whether supercoiling changes cleavage positions within off-targets, despite recent structural support^35^. This is especially critical, as changes in cleavage position alter editing outcomes with Cas9^41^.

In this high-throughput study, we investigate how supercoiling impacts where and how quickly Cas9 cleaves off-targets. Our results offer genome-agnostic, nucleotide-resolution kinetics and DNA end analysis for supercoiled libraries of systematically-mispaired off-targets. By expanding application of the NucleaSeq method^15^ to supercoiled targets, we directly relate our results to our previous study on relaxed DNA. We show that Cas9’s ability to cleave targets depends on the mispairs present and the target’s topology. Supercoiling increases Cas9’s cleavage kinetics for off-targets with seed-region mispairs, but cleaves off-targets with PAM-distal mispairs similarly whether supercoiled or not. We observe that active Cas9 either nicks or repositions cleavage of some off-targets with PAM-proximal indel mispairs. Finally, we use our biophysical CRISPRzip^42^ model to rationalize the effects we observed in supercoiled target experiments. The model allows us to predict Cas9 cleavage kinetics across target sequences and supercoiling levels. Together, our data and model show the range of Cas9 activity users can expect across the DNA topologies found in our genome.

## Results

### NucleaSeq captures Cas9 cleavage and binding kinetics on supercoiled DNA

The NucleaSeq method measures nuclease cleavage kinetics and cleavage position for relaxed, linear DNAs^15^. However, negative supercoiling (nSC) plays an important role in Cas9 targeting specificity^17,33,34^, so we adapted NucleaSeq to capture its effect on nuclease specificity during cleavage (Figure 1A). Our libraries^15,43^ included targets that perfectly matched Cas9’s PAM and gRNA sequences gRNA1 and gRNA2 (as used in^15,22,34,44–47^) those with alternative PAMs, non-target control sequences, and thousands of off-target sequences with gRNA-relative single or double mismatches, insertions and deletions (Figure 1B). Each target sequence was flanked by unique barcodes for post-cleavage identification.

**Figure 1.**
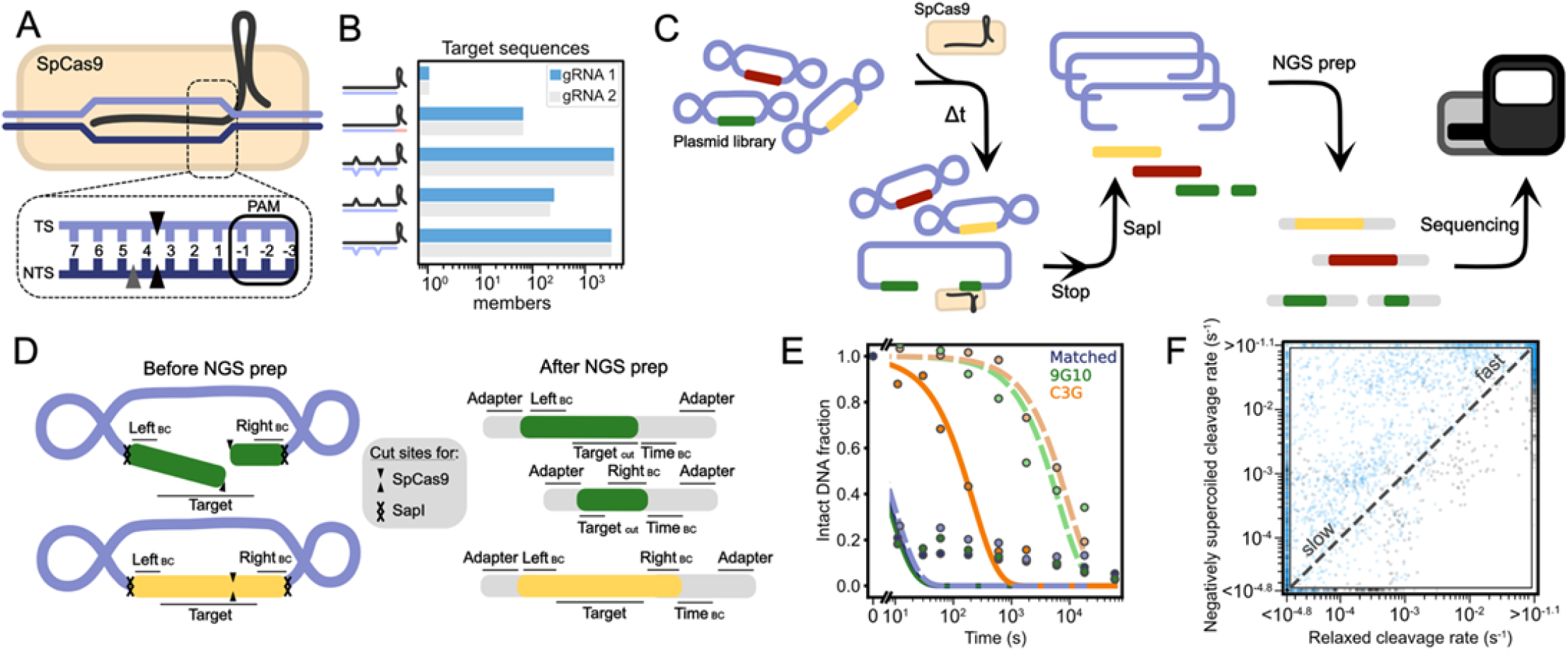
An adapted NucleaSeq method captures cleavage rates and sites for targets in supercoiled plasmid DNA. (A)Cas9 typically cleaves on-target DNA between the 3^rd^ and 4^th^ bases of the target (TS) and non-target (NTS) strands (black arrows). Some on-targets experience non-target strand (NTS) cleavage between the 4^th^ and 5^th^ bases (grey arrow). (B)Composition of targets in plasmid libraries for gRNA 1 and 2. Both libraries include fully-matched (gRNA and PAM) on-targets, off-targets with alternative PAMs, and off-targets with one or two mismatches, insertions and deletions relative to the gRNA. (C)The updated NucleaSeq workflow assays plasmid cleavage. Plasmid library is exposed to Cas9 for increasing times (Δt = 0, 0.16, 0.5, 1, 3, 10, 30, 100, 300, 1000 min) before quenching (Stop). The quenched reactions contain cleaved and intact targets. Then, samples are restricted with SapI and the plasmid vector is removed from library targets based on size. DNAs remaining in each sample receive a time barcode during NGS preparation, and are finally sequenced by short-read NGS. (D)A diagram shows individual cut (green) and intact (yellow) DNAs from a supercoiled target library before (left) and after (right) NGS prep. Initially, two unique barcodes (Left BC, Right BC) and SapI sites (hatch) flank each target. Cas9 cleavage (arrows indicate site) separate one side’s unique barcode and SapI site from the other side’s. Whether cut or intact, each piece of DNA receives a time barcode (Time BC) during the adapter ligation step of NGS preparation. (E)Fraction of DNA remaining intact upon Cas9 exposure over time for gRNA and PAM-matched on-target (Matched, purple) and two off-targets (9G10, green; C3G, orange). Read counts (circles; intense colors=nSC while matched pastel colors=Rel) are normalized at each time point to non-target controls, and then to the t=0 read counts. Exponential functions are fit to read normalized counts from supercoiled DNA (solid lines) or relaxed DNA (dashed lines). (F)Library-wide comparisons of cleavage rates for supercoiled versus relaxed target DNA. Blue: Library 1 for gRNA 1. Grey: Library 2 for gRNA 2. Dashed line shows x=y.

We inserted each of our libraries’ members into a minimal 1.6kb plasmid vector and transformed them into *E. coli*. Purified plasmid libraries were negatively supercoiled, with a superhelical density of σ=-0.033±0.007 (Supp. Fig. 1) ^48^. For nSC cleavage kinetics via NucleaSeq, we exposed each plasmid library to the Cas9 ribonucleoprotein complex (RNP) at 22°C, periodically quenching samples from the reaction (see methods). After removing the plasmid vector (Figure 1D), we appended time-associated barcodes (*e*.*g*. sequencing indices) during preparation for next generation sequencing (NGS). Samples were pooled together and sequenced (Figure 1C). Sequencing read counts for each library member and timepoint were normalized (as described in ^15^) and fit to exponential functions to determine effective cleavage rates (Figure 1F).

### Supercoiling alters Cas9 off-target cleavage rates and positions

To investigate Cas9 specificity across targets when cleaving nSC DNA, we investigated off-targets with three mispair types – mismatches, insertions or deletions (when comparing target DNA to the gRNA used, Figure 2). Mismatches can involve non-canonical base pairing, while insertions and deletions can introduce nucleotide bulges and base skipping^49^. Our previous NucleaSeq results used the same libraries and analysis as present^15^, enabling direct comparisons between relaxed and nSC DNA cleavage kinetics. Relaxed DNA off-targets experience very slow cleavage by Cas9 when containing PAM-proximal “seed” mispairs as opposed to PAM-distal mispairs (Figure 2A). Here, Cas9 cleaved these same off-targets much faster when negatively supercoiled, regardless of whether mispairs were proximal or distal, in agreement with a prior study^39^, and across our target libraries (Fig 2A). Cas9 cleaved nSC off-targets with single mismatches in the seed region up to 100-fold faster than relaxed counterparts, while rates for most PAM-distal nSC off-targets were so fast as to be indistinguishable from the on-target rate. Surprisingly, off-targets with a single deletion mispair in the first 4 PAM-proximal positions were cleaved slower when supercoiled than relaxed. Such unpaired bases may misalign the target DNA with Cas9’s cleavage domains, and supercoiling could exacerbate this misalignment^49^. Off-targets with insertions at positions 6-10 showed much faster cleavage on nSC DNA than relaxed DNA, but not so for off-targets with insertions at positions 11-17 – a region where hybridization licenses Cas9 cleavage^50^. Among single mispair off-targets, the difference between nSC and relaxed DNA cleavage rates was greatest for insertion mispairs: some were cleaved >10k-fold faster when supercoiled. These results highlight that Cas9 effectively ‘queries’ DNA topology when discriminating off-targets.

**Figure 2.**
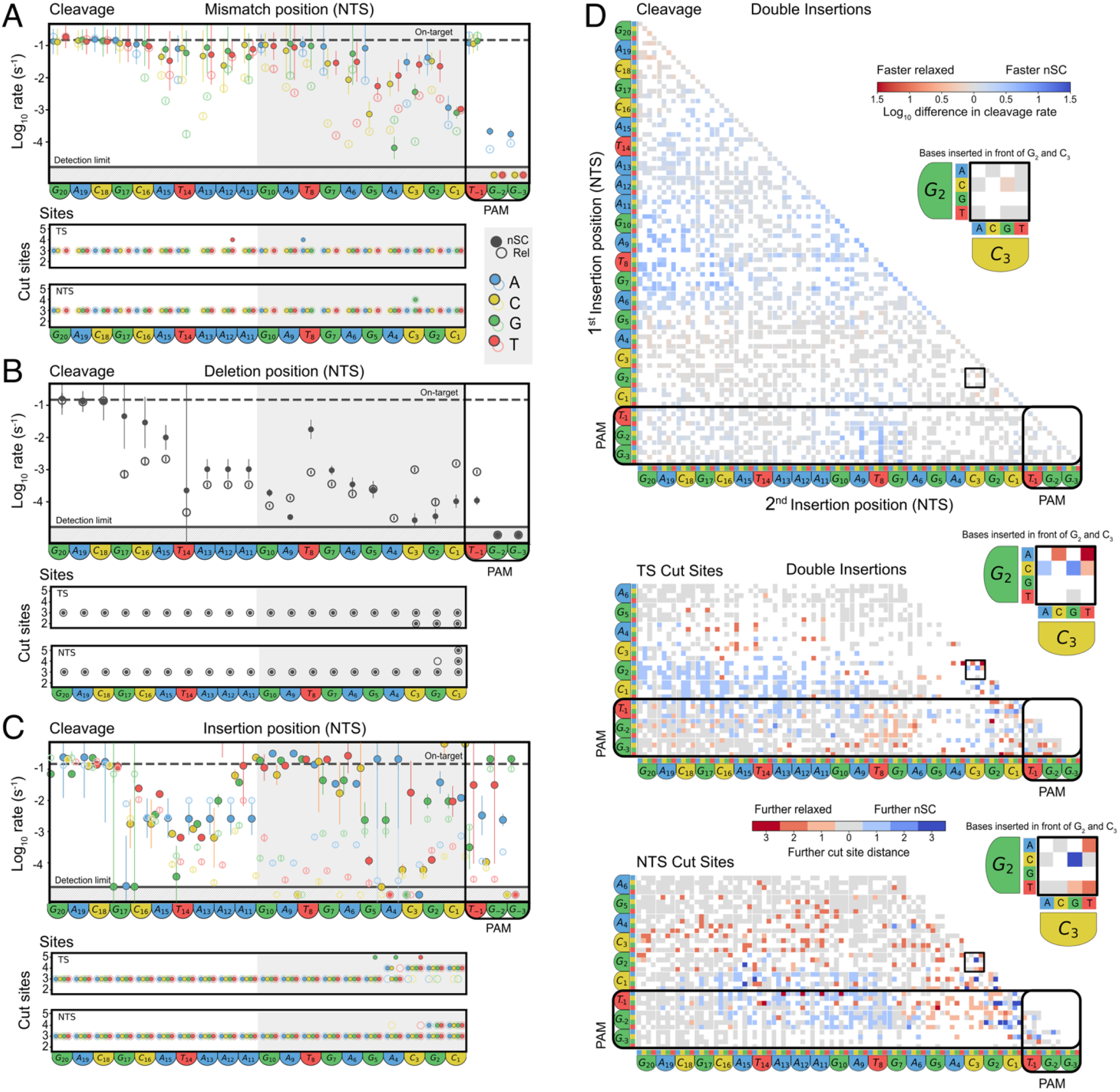
Supercoiling affects Cas9 binding and cleavage rates as well as position of cleavage in a mispair-dependent manner. Fitted Cas9 cleavage rates and cleavage positions for targets with a single PAM- or gRNA1-relative mismatch **(A)**, deletion **(B)**, or insertion **(C)**. Hollow circles: relaxed target values. Filled circles: supercoiled target values. For cleavage graphs, targets with fewer than 30 reads at t=0 were omitted. Error bars: SD based on 50 rounds of bootstrapping with replacement. X axis labels: NTS nucleotides and positions in relation to the PAM and gRNA1. Grey box: approximate seed region. Dashed lines: on-target cleavage rate (nSC and Rel). Solid lines: detection limits. Hatched regions contain fit values exceeding the adjacent detection limit. Target cleavage positions are shown for the target strand (TS, top) and non-target strand (NTS, bottom). Indicated cleavage positions were found in ≥10% of cut reads for one or more timepoints. Cleavage positions are calculated away from the PAM. X axis labels: NTS nucleotides and positions in relation to the PAM and gRNA1. **(D)** Epistasis of Cas9 cleavage rates and cleavage positions for nSC and relaxed DNA with two PAM- or gRNA-relative insertions. X and Y axes show the NTS sequence, and the x:y intersection shows the difference in cleavage rate or the further cut site for the double mispair. For cleavage data, intensifying shades of blue show a faster cleavage rate on nSC compared to the relaxed DNA, while intensifying red shades show faster relaxed DNA cleavage compared to nSC. For cut sites, shades of blue correspond to the number of nucleotides nSC DNA was cut further away than corresponding relaxed DNA and shades of red correspond to the number of nucleotides relaxed DNA was cut further away than corresponding nSC DNA. For both graphs white: omitted targets with fewer than 30 reads at t=0, grey: no difference between nSC and relaxed cleavage rate or site. Black boxes expanded in callouts.

We next examined off-targets containing two mispairs (Fig 2B). As with relaxed off-targets, Cas9 could still cleave some nSC off-targets containing multiple mispairs, in agreement with cell-based studies (Supp. Table 1) ^20^. As with single mispairs, double-mispair off-targets were often cleaved faster when negatively supercoiled (Figure 2D). This was most apparent for off-targets with an insertion mispair in positions 6-10, except when preceded by another insertion mispair near the PAM (positions 2-4). For off-targets with two deletion mispairs, Cas9 cleaved them so slowly that topology was irrelevant. These results indicate that the impact of supercoiling and mispairing are not simply additive.

Finally, we analyzed differences in Cas9 cleavage sites library-wide, with respect to supercoiling. Cas9 typically cleaves on-target DNAs between the third and fourth nucleotides from the PAM, or occasionally the fourth and fifth on the non-target strand (depending on the gRNA)^3,15^. Similarly, mispairing and supercoiling might alter the orientation of each DNA strand to its nuclease domain, thereby shifting cleavage sites. We found that nSC off-targets with single mismatches were largely cleaved at the same positions as their relaxed counterparts, and similarly to the on-target (Figure 2C)^15^. However, for off-targets with single PAM-proximal deletions, Cas9 often cleaved one nucleotide closer (on the target strand; TS) or further (on the non-target strand; NTS) from the PAM, independent of supercoiling. Off-targets with single PAM-proximal insertions showed the greatest topology sensitivity: upon supercoiling, target strand cleavage shifted as much as two nucleotides further from the PAM. This may result from downstream R-loop hybridization more strongly impacting nucleotide positioning than PAM-PID interactions.

We investigated whether shifts in cleavage sites extended to off-targets containing two mispairs (Figure 2D). Off-targets with two PAM-proximal insertions were often cleaved two or more nucleotides further from the PAM (Supp.Table 1) on the TS, regardless of topology. However, off-targets with a PAM-proximal (positions -1 – 3) and a PAM-distal (∼10+) insertion mispair showed a greater shift in cleavage site for the TS of nSC off-targets. This contrasted with NTS, where Cas9 cleaved further away on relaxed off-targets than on nSC off-targets. Thus, while Cas9 slowly cleaves off-targets with PAM-proximal mispairs, these off-targets nonetheless experience the greatest shift in cleavage rates and sites depending on DNA topology.

### Cleavage site-adjacent mispairs disrupt Cas9 nuclease activity

Mispairing between the gRNA and target strand DNA distorts the heteroduplex structure within Cas9^51^. Supercoiled off-targets (with PAM-proximal insertion mispairs) can experience Cas9 cleavage further from the PAM (Figure 2C) indicating that supercoiling further distorts the heteroduplex. We hypothesized that this distortion impacts cleavage by Cas9’s RuvC and HNH nuclease domains differently, which cannot be detected via dsDNA cleavage alone. We selected three off-targets for deeper investigation: ‘C3G’, ‘Δ3’, and ‘3A4’ (DNA with G-C replacing C-G at position 3, no position 3 base pair, or an extra A-T between base pairs 3 and 4, with respect to the gRNA-matched DNA sequence) (Figure 2B). For Δ3 and 3A4, Cas9 nicked – but did not cut – the DNA (Figure 3A). This suggests that at least one nuclease domain loses its cleavage capacity where “extra” DNA or gRNA bases distort local pi-stacking and structure, a phenomenon not observed with just a single mismatch^52^ (Figure 3A).

**Figure 3.**
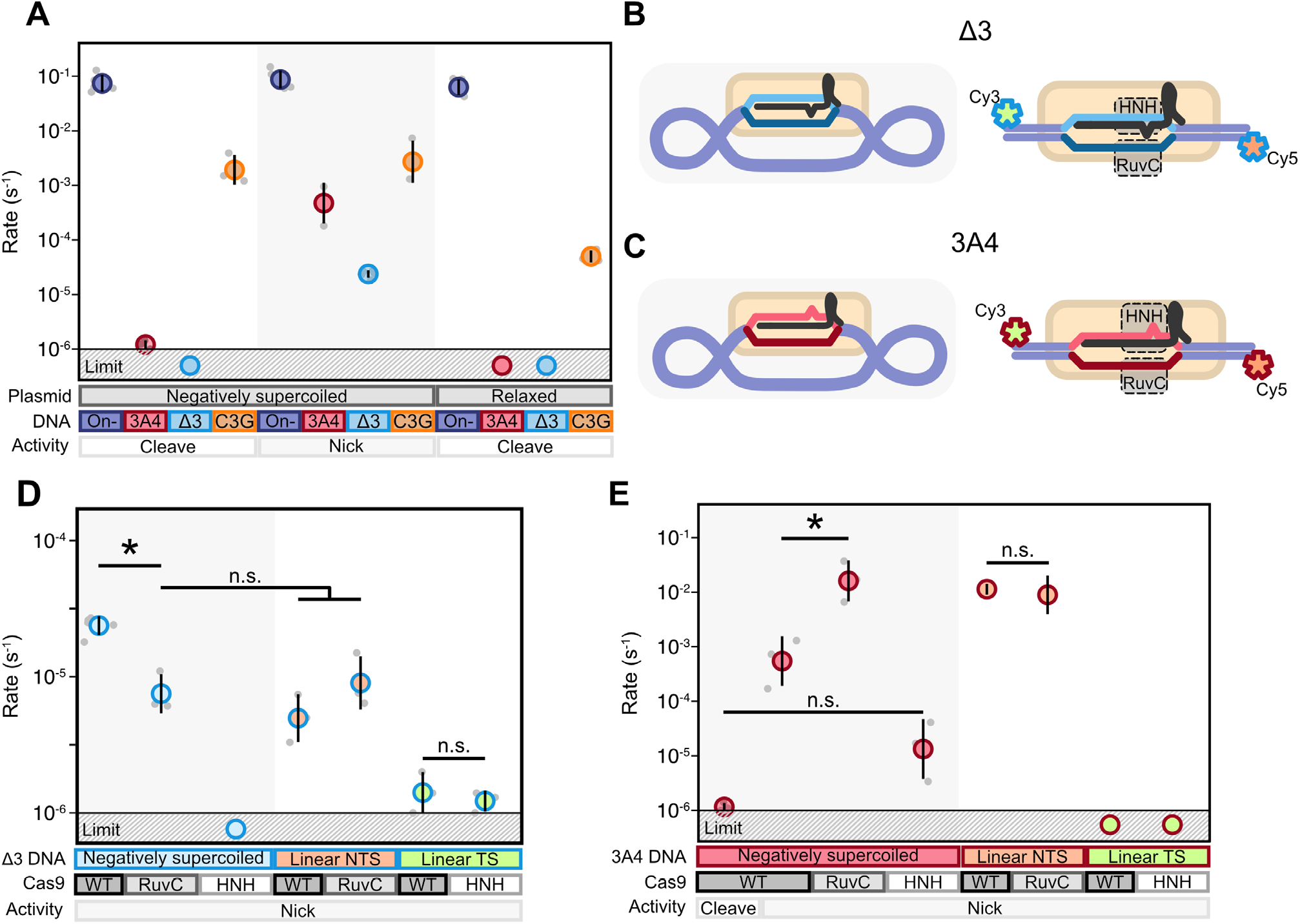
Cleavage site-adjacent mispairing differentially inhibits Cas9 nuclease domains. **(A)** Cas9 cleavage and nicking rates of supercoiled or relaxed plasmids containing off-targets with mispairs at position 3. Single exponential functions were fit to lane-normalized agarose gel band intensities for all samples with detectable cut DNA in at least the last two timepoints (300min and ≥1000min). Solid line: detection limit based on maximum ethidium bromide sensitivity on the gel. Mean ±SD. N≥3 for all targets. Schematics for **(B)** off-target Δ3 DNAs used in D, and (C) off-target 3A4 DNAs used in E. **(D)** Fit nicking rates for off-target Δ3 (gRNA1-relative deletion) when present in supercoiled plasmid DNA or relaxed linear DNA. WT Cas9, ΔH840A Cas9 nickase (active RuvC) or D10A Cas9 nickase (active HNH) were used as indicated. Single exponential functions were fit to lane-normalized agarose gel band intensities for all samples with detectable cut DNA in at least the last two timepoints (300min and 1000min). Relaxed linear DNAs were co-labeled with a Cy5 fluorophore on the non-target strand (NTS) and a Cy3 fluorophore on the target strand (TS). Solid line: detection limit based on maximum ethidium bromide sensitivity on the gel. Mean ±SD. N≥3. * p ≤0.05 with Tukey’s post-hoc test, after testing for normality. **(E)** Fit cleavage or nicking rates for off-target 3A4 (gRNA1-relative insertion) when present in supercoiled plasmid DNA or relaxed linear DNA. WT Cas9, ΔH840A Cas9 nickase (active RuvC indicated) and D10A Cas9 nickase (active HNH indicated) were used. Single exponential functions were fit to lane-normalized agarose gel band intensities for all samples with detectable cut DNA in at least the last two timepoints (300min and 1000min). Relaxed linear DNAs were individually labeled with a Cy5 fluorophore on the non-target strand (NTS) or a Cy3 fluorophore on the target strand (TS). Solid line: detection limit based on maximum ethidium bromide sensitivity on the gel. Mean±SD. N≥3. * p ≤0.05 with Tukey’s post-hoc test, after testing for normality.

To pinpoint which Cas9 cleavage domain becomes impaired – HNH or RuvC, we quantified single and double nicking events by WT Cas9 and HNH- or RuvC-active only Cas9 nickases on supercoiled Δ3 and 3A4 off-targets (Figure 3B-E). Then, we measured HNH and RuvC behavior individually by performing these experiments with relaxed, fluorescently-labeled DNAs (Figure 3B-E).

Wildtype Cas9 (WT Cas9) only nicked nSC Δ3 (Figure 3B) off-target plasmid once, with no detectible second nick activity (Figure 3D). When investigating this nSC plasmid with nickase RNPs, only the RuvC-active Cas9 generated a nick. This suggested that WT Cas9’s RuvC operates on this target, but its HNH does not. When investigating Cas9 activity with fluorescently labelled DNA (Figure 3B), both domains generated cleavage products, albeit only faintly for HNH (Figure 3D). This activity agrees with Cas9’s known topology-independent nicking activity^39,52,53^. When investigating our off-target with an unpaired gRNA nucleotide at the typical cleavage site (target 3A4, Figure 3C), we observed topology-dependent nicking: HNH activity only occurred when the target was supercoiled (Figure 3E). Thus, we report for the first time that a target’s sequence *and* topological state together convert Cas9 from a nuclease to a nickase with as little as a single mispair between the gRNA and target.

### A biophysical model captures the impact of supercoiling on Cas9 specificity

To obtain physical insight into our observed effects of supercoiling on Cas9 specificity (Figures 1-2), we applied our mechanistic ‘CRISPRzip’ model^42^. CRISPRzip incorporates the effects of torque on Cas9 activity as previously demonstrated on single molecules^34^. This model leverages the fact that Cas9’s R-loop forms progressively during target interrogation (Figure 4A). CRISPRzip simulates the kinetics of this process as stochastic progress through sequence-dependent free-energy landscapes (Figure 4B, C). We obtained the underlying landscape parameters by training on (d)Cas9 (binding) cleavage dynamics for relaxed targets with up to 2 point mutations (library-1, Supp. Fig. 2). ^42^ We integrated the effect of torque^34^ by letting DNA underwinding reduce the work needed to extend the R-loop, effectively tilting R-loop landscapes downward and accelerating cleavage (Figure 4B, C). DNA overwinding does the opposite.

**Figure 4.**
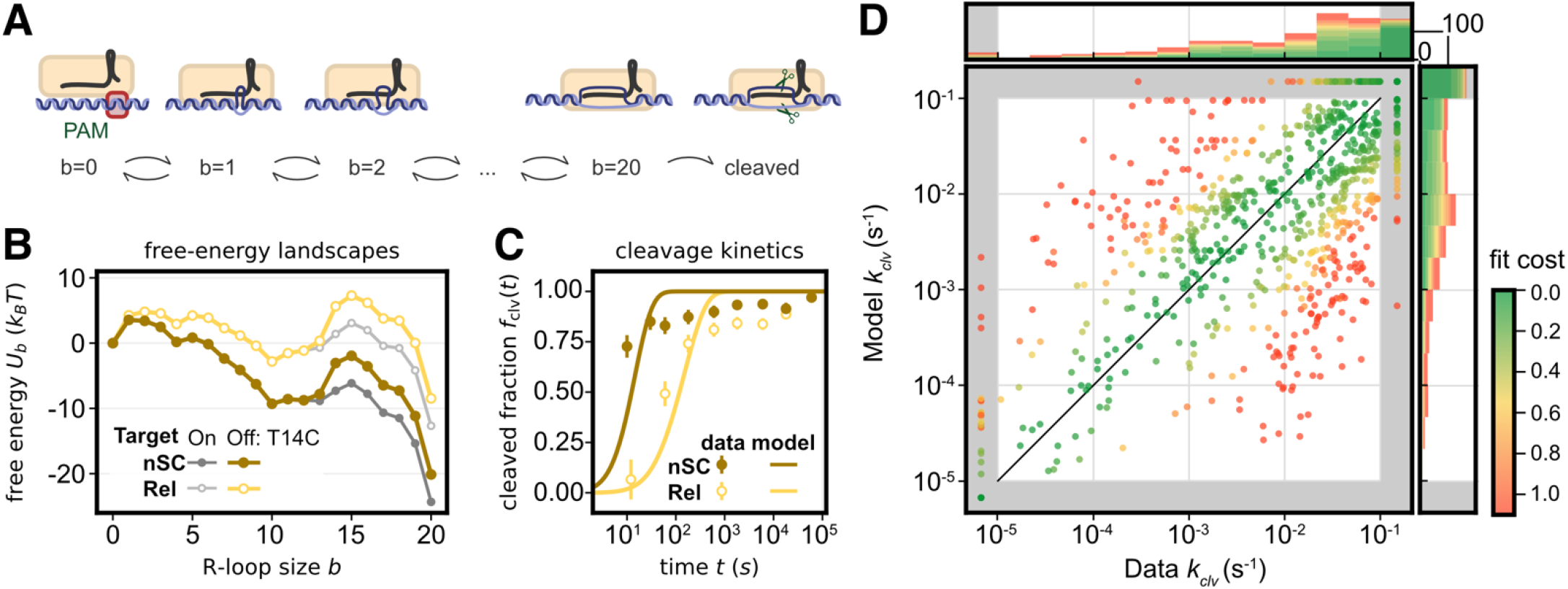
Modelling the mechanistic effects of supercoiling. **(A)** The CRISPRzip model simulates the kinetics of R-loop formation at base pair resolution. At state *b*, where *b* hybrid DNA:RNA base pairs are established, the free-energy difference with the neighboring states *b* ™ 1 and *b* + 1 respectively determine the rates of R-loop extension and recession. **(B)** Modeled free-energy landscapes of R-loop formation for off-target T14C DNA when relaxed (yellow) or supercoiled (brown). Free-energy landscapes for the on-target are shown for reference (grey). Negative supercoiling effectively tilts the landscapes down. **(C)** Cleavage of target T14C, as measured experimentally (circles) and as predicted with CRISPRzip (solid lines). The negative landscape tilt results in accelerated cleavage. Error bars: SD based on propagated uncertainty from Poisson-distributed initial read counts. **(D)** Measured vs CRISPRzip-predicated cleavage rates *k*_clv_, after training CRISPRzip on the library-1 nSC cleavage data, which resulted in a fitted plasmid torque of τ_0_ = − 3.6 ± 1.1 pNnm. Circles: data points for targets from library 1. Colors indicate fit cost of that target.

To assess if our high-throughput kinetic data can be explained mechanistically, we trained the torque level in CRISPRzip to the cleavage kinetics measured across library 1, as it contained the largest target set (Figure 1B). The obtained estimate for the plasmid torque (τ_0_ = −3.6 ± 1.1 pNnm) corresponds to plasmids with superhelical density σ = −0.03 ± 0.01^42^, which agrees well with our supercoiling estimates from gel electrophoresis (Supp.Fig. 1). Visualizing the predicted and measured cleavage rates shows that CRISPRzip reproduces the cleavage kinetics of most targets, spanning the full dynamic range (Figure 4D). The remaining variation may be due to sequence-specific responses to supercoiling that are outside the model scope.

We next leveraged CRISPRzip to understand the relationship between Cas9 off-target binding and cleavage upon increased negative supercoiling. While binding is necessary for cleavage to occur, even transient binding suffices. Weakly-bound off-targets may be cleaved, just as well-bound targets may not be^42^ (Figure 2A,B). CRISPRzip modeling indicates that ‘fast-cleaved but weakly-bound’ targets become more stably bound with increased negative supercoiling (Figure 5A; targets in the top left clear sooner upon nSC). Further, CRISPRzip predicts the impact that positive supercoiling has on Cas9 activity^42^ – as seen during replication, transcription and in archaeal genomes (Figure 5B).

**Figure 5:**
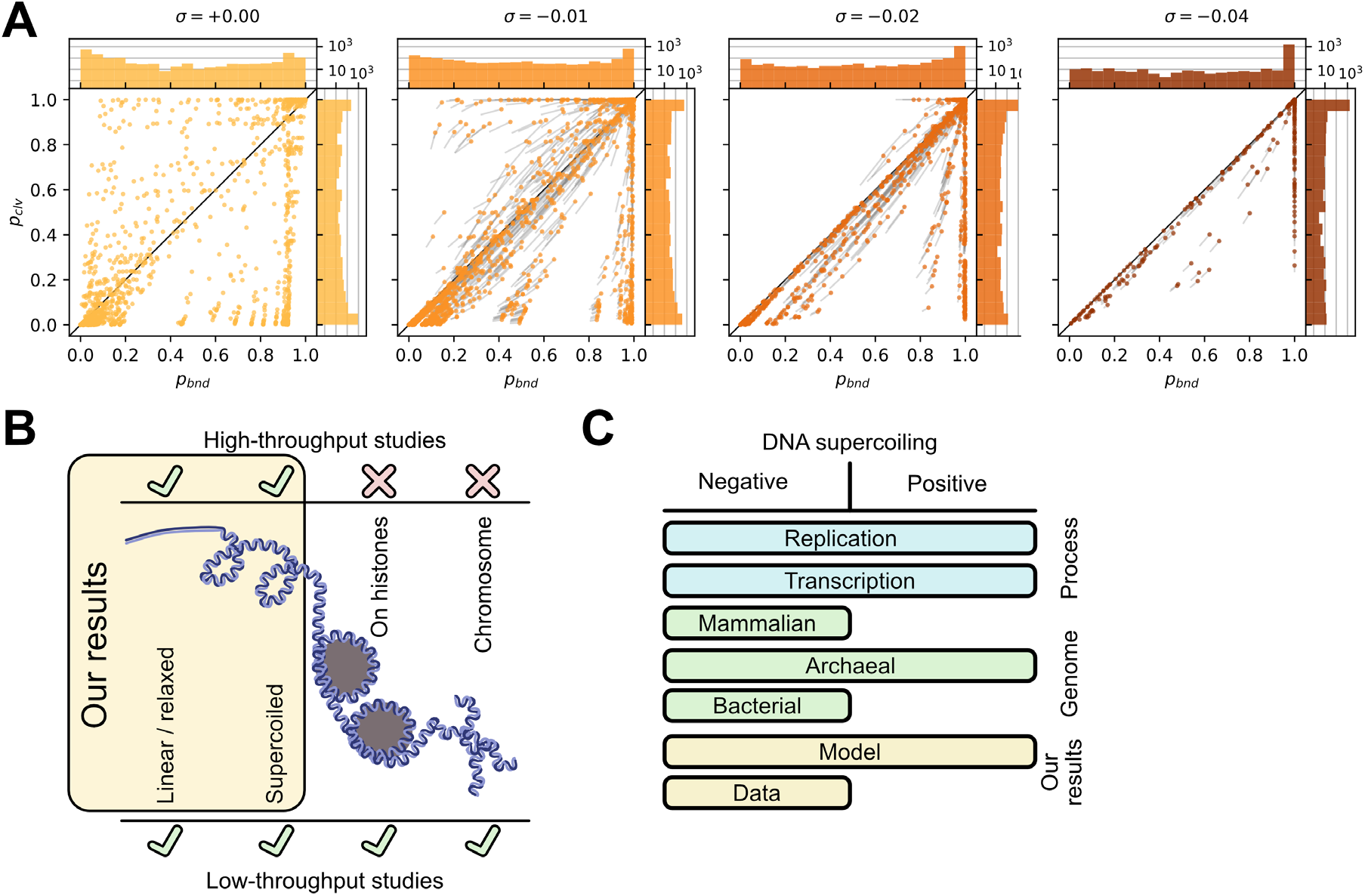
CRISPRzip predicts the mispair-dependent role of supercoiling in Cas9 binding and cleavage. **(A)** CRISPRzip predicts the probability that dCas9 binds (*p*_*bnd*_) and Cas9 cleaves (*p*_*clv*_) all targets with 0-2 mismatches after 15 minutes of exposure to 20 nM RNP. Tails (gray) show each target’s sensitivity to supercoiling as changes in *p*_*bnd*_ and *p*_*clv*_ over σ +0.001 to σ, where σ = superhelical density. **(B)** CRISPRzip predicts Cas9 activity for negative and positive DNA supercoiling, as found across genomes and cellular processes. **(C)** High-throughput *in vitro* studies provide greater resolution and context to results from lower-throughput studies that describe how DNA organization impacts Cas9 specificity and utility for genome editing.

## Discussion

Here, we investigated Cas9 off-target discrimination in a genome-agnostic manner, exposing its intrinsic binding and cleavage specificity, and its dependence on target supercoiling. We used the biophysical model CRISPRzip for a kinetic understanding of (d)Cas9’s supercoiling sensitivity. Our work complements prior studies, which typically investigate Cas9 specificity in cellular environments or on extracted genomes (human, mouse, *etc*.)^54,55^. Such results capture the impact of unique genome characteristics like chromatin accessibility and transient topologies on Cas9 specificity. Thus, off-target models built on these results exchange robust application across genomes for accuracy within the genome used for training^54,56^. Indeed, differential gene expression across cell cycle phases and types dramatically impacts editing efficiency, editing outcomes and off-target activity ^55,57,58^. Our use of synthesized libraries profiles Cas9 specificity with challenging yet comprehensive target sets that exceed any individual genome’s contents.

Supercoiling represents a basic, nearly-universal aspect of topological DNA organization, yet is the highest topological level for which high-throughput biochemical Cas9 specificity studies are available (Figure 5C). The next step is compaction upon nucleosomes. Available *in vitro* studies have examined extremely tight nucleosome wrapping, which effectively prevents Cas9 access ^59–61^. However, natural nucleosome breathing (and chromatin remodeling) reveals targets, where cleavage rates become equivalent to naked DNA targets^60,62^. Further, Cas9 dwells on cellular targets for several minutes after cleavage^63^ but experiences dislocation by other factors^64^, thereby operating in a multiple-turnover regime. Thus, understanding how nucleosome positioning and movement impacts Cas9 activity and specificity remains a critical, but largely under-defined, factor that limits predictable action in the cell.

Target-gRNA mispair tolerance is also observed across other CRISPR-Cas enzymes: CRISPR-Cas12a cleaves off-targets with mispairs and non-canonical PAMs^15,65^, as does Cas12b, Cas12j, and Cas9 variants. For Cas12a, R-loop propagation and cleavage rates across off-targets strongly correlate, as its Cas12a’s mechanism relies heavily on full R-loop propagation for successful dsDNA cleavage^66^. Even so, Cas12a shows promiscuous and accelerated cleavage for supercoiled targets^67–69^, while Cascade shows very low binding *and* cleavage activity on relaxed targets^70–72^. Within the Cas9 family, archaeal *At*Cas9 and *Ah*Cas9, supercoiling nearly abolishes PAM requirements for supercoiled targets, but not for relaxed ones^73^. Enzymes from other defense systems, like Gabija, also show preference for supercoiled DNA for its cleavage^74^. Thus, Cas9’s mispair tolerance reflects a typical, rather than exceptional, feature found in defense systems.

Our results show two “proofs-of-concept” where Cas9 could be developed: as a supercoiling detector, or a mutation-free nickase. We report here that some off-targets experience a substantial increase in Cas9 cleavage rate when negatively supercoiled (*e*.*g*. 9G10, Figure 2B, Supp. Fig. 3). By programming Cas9 with a gRNA that recreates such a mispair, one would effectively convert DNA cleavage into a sensor for DNA torque. This could offer a simple and quick alternative to intercalator assays^48^. We also reported for the first time that just a single-base mispairing (3A4, Δ3; Figure 3D,F) can suppress HNH domain activity, converting WT Cas9 into a nickase. Base and prime editing are both enhanced by nicking, where prime editing uses HNH-inactive Cas9^75,76^. Thus, mispairing gRNAs may offer reinforcement, or potential replacement, of inactivating mutations in such applications.

## Materials and Methods

### gRNAs

All gRNAs used in this study were purchased from Synthego or GenScript and diluted in 1 mM Tris, 0.1 mM EDTA, pH 8.0 (Table 1).

**Table 1.**
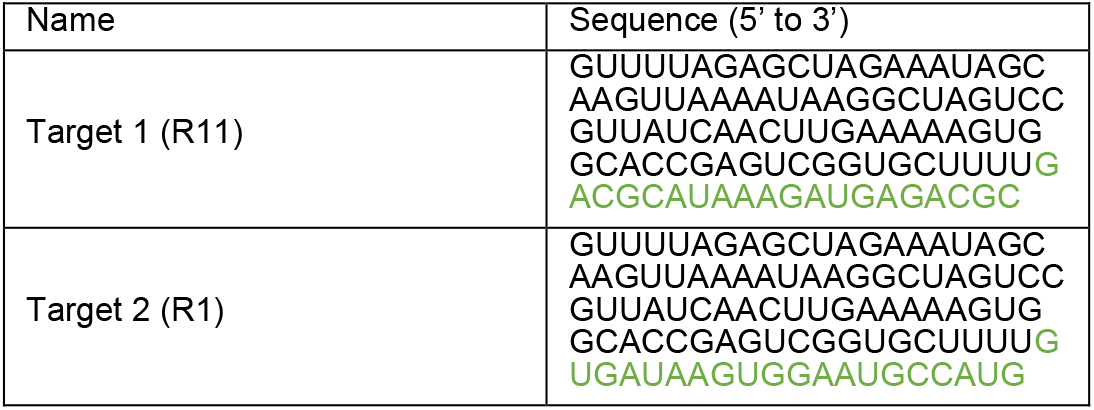
gRNAs used in this study (target sequence in green, scaffold in black):

### Plasmid library construction

This study used barcoded target libraries (GenScript)^15,43^ amplified by PCR (Phusion Plus polymerase, Thermo Scientific, Table 2, primers Pr6.29/Pr6.30 for library 1; Pr7.26/Pr7.27 for library 2) with 25 cycles or less to maintain library diversity and then column purified (GeneJET PCR purification kit, Thermo Scientific). Plasmid backbone (Table 2, Pl0.63) was similarly amplified with 20 cycles or less with Pr6.31/Pr6.32 for library 1 and Pr7.28/Pr7.29 for library 2 (Table 2), then column purified. Both libraries were inserted into empty plasmid backbone via HiFi assembly (NEB), using a 5:1 ratio of library to backbone.

**Table 2.**
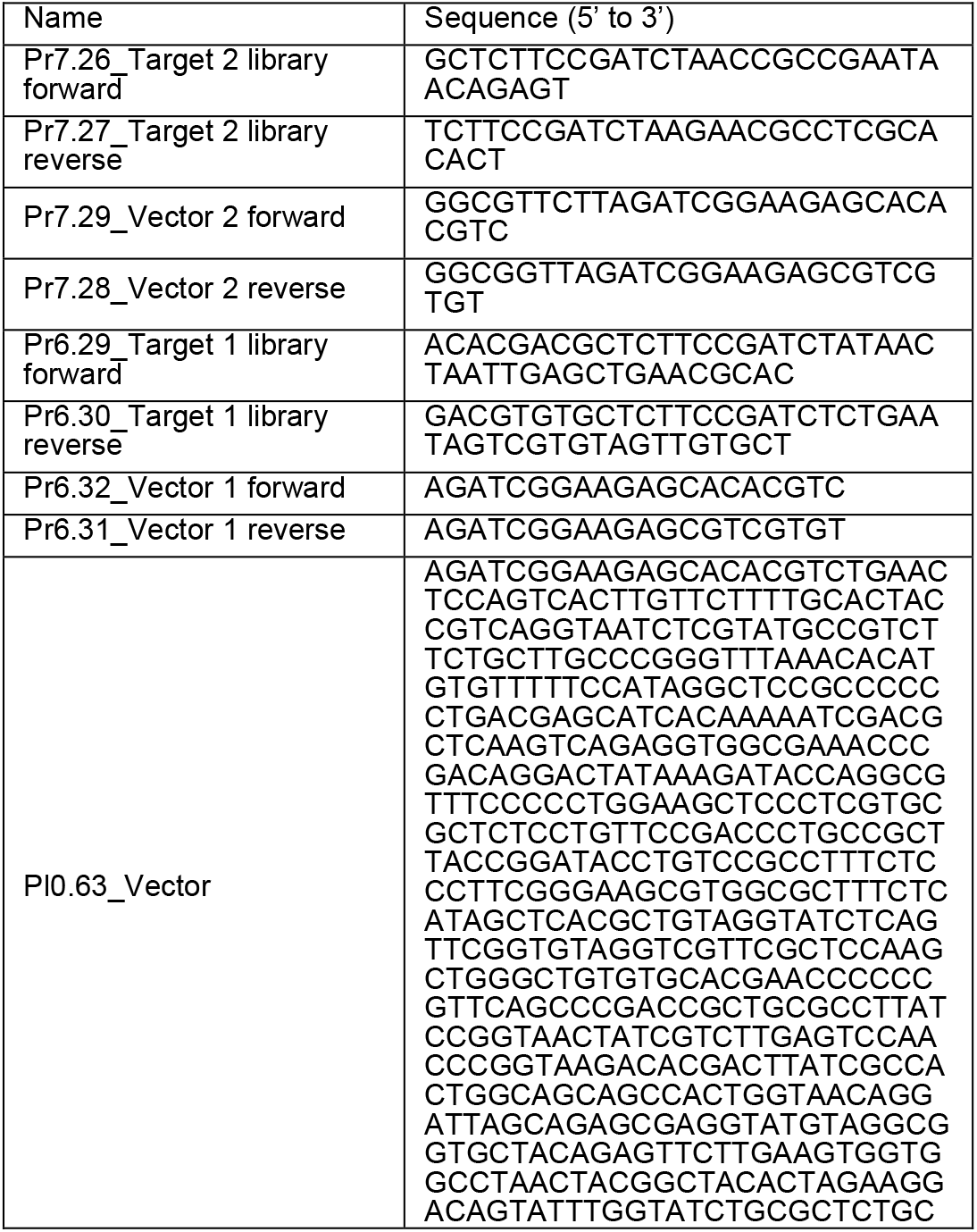
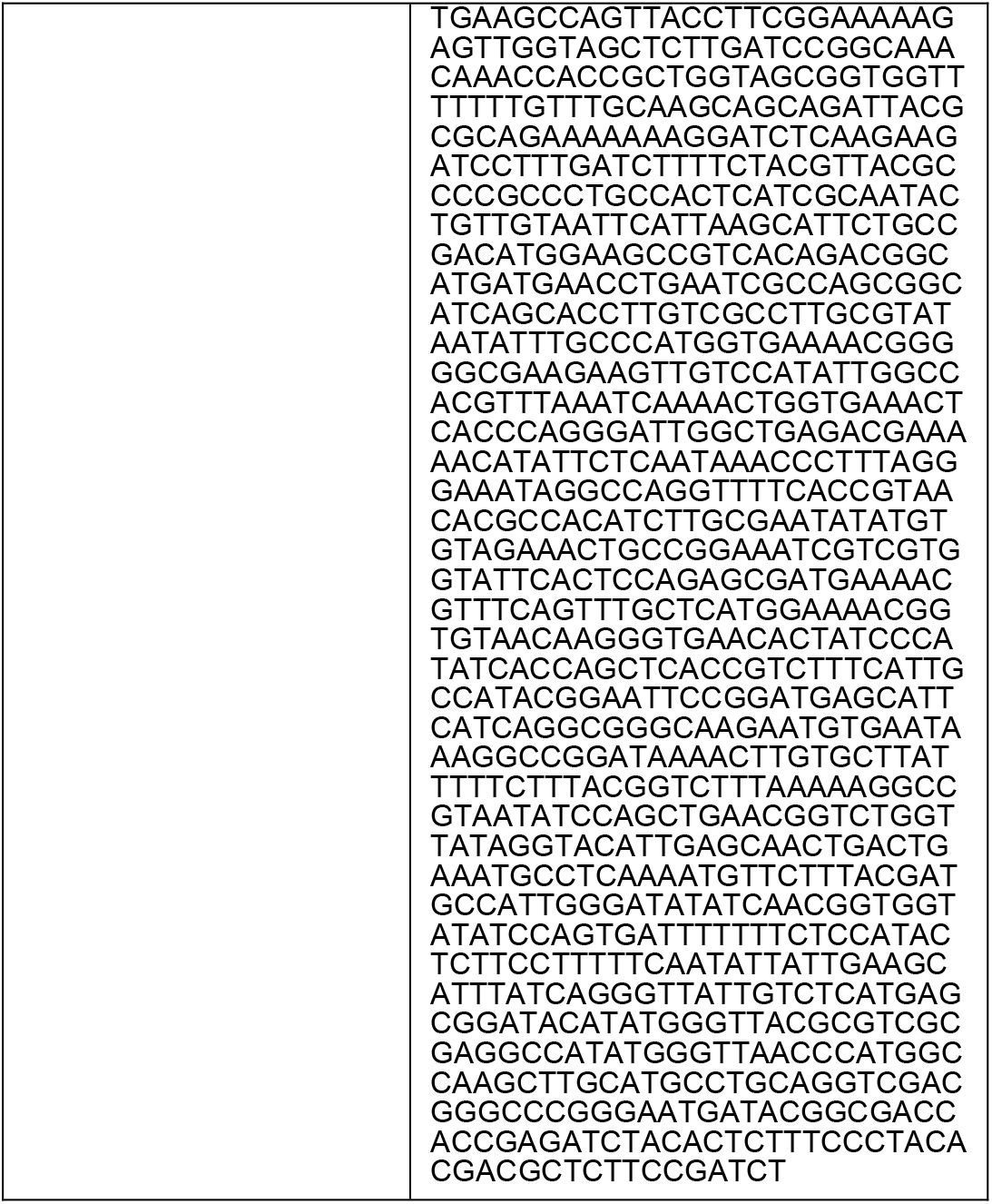
Primers and plasmid backbone used (from Metabion or Azenta):

### Plasmid library transformation and extraction

HiFi-assembled libraries were electroporated (ECM630, BTX; 1.8kV) into TOP10 E.coli cells (Invitrogen). To ensure efficient transformation and sufficient library representation, 10^4^-fold dilutions were plated on Lb Agar + chloramphenicol (25μg/mL) and grown at 37°C for 20h. Target libraries 1 and 2 had ≥120 and ≥90 colonies per library member collected from one or more transformations. Transformed cells were plated on 3+ 150mm Lb agar plates with chloramphenicol (25μg/mL). After growing the cells at 30°C for 20-22h, the plates were scraped with 3mL LB media a total of three times, and then purified (Pure II Plasmid Midiprep kit, Zymo Research, D4201).

### Superhelical density determination

To determine the superhelical density of purified plasmids, a previous protocol was modified^48^ to accommodate our compact plasmid (1.7 kb). A subset of purified plasmids was relaxed with Topoisomerase I (NEB). Supercoiled and relaxed plasmids (0.5μg each) were ran on a 1% agarose gel (TBE, 2.4μg/mL of chloroquine diphosphate (Fisher Scientific) in the gel and the buffer) at 3V/cm for 4 hours, then post-stained with SYBR Safe (Invitrogen). We calculated band intensity profiles using GelAnalyzer 19.1. The change in linking number distribution (ΔLk) was calculated by counting bands from the middle of relaxed DNA to the middle of supercoiled DNA bands (Supp. Fig. 1), and then superhelical density was calculated as described in ^48^.

### NucleaSeq

The reaction components – sgRNA (Table 1), Cas9 (NEB), plasmid library – were diluted in 100 mM NaCl, 50 mM Tris-HCl, 10 mM MgCl_2_, 100 µg mL^−1^ recombinant albumin, pH=7.9 (r3.1 buffer, NEB), to the final reaction concentrations of 187.5nM, 62.5nM, 6.25nM, respectively. First, diluted sgRNA and Cas9 were mixed to form an RNP for 15 minutes at 22°C. After the addition of the plasmid library, small aliquots of the reaction were removed at different timepoints – 0, 0.2, 0.5, 1, 3, 10, 30, 100, 300 and 1000 min – and added to the STOP solution (60mM EDTA (Carl Roth), 1U Proteinase K (Thermo Scientific)), then incubated at 37 °C for 30 minutes.

Timepoint 0 was treated with STOP solution before the addition of DNA. Each timepoint sample was then ethanol precipitated.

The cleaned timepoint samples had their backbones cut out with an excess of SapI (NEB). The resulting reactions were measured with High Sensitivity Assay (Qubit) and half of each reaction was used for appending time indexes with NGS prep (NEBNext, NEB). Barcoded samples were size-selected with AMPure XP magnetic beads (Beckman Coulter) using 0.9X bead:DNA ratio, ran on capillary electrophoresis (BioAnalyzer 2100, Agilent) and then sequenced on a NextSeq2000 (P2 100bp PE or P3 100bp PE chemistry) at the EMBL GeneCore (Heidelberg, DE) with 10% PhiX spike to increase sequence diversity.

### NGS data analysis: identification, normalization and rate fitting

Read counts for each library member were tabulated as previously described^15^. Briefly, paired NGS reads were merged, then filtered by quality and size. Acceptable reads for intact DNAs required correct matching to both barcodes and the target for a single library member, while cut DNAs required correct matching to a single barcode and the portion of the remaining target. After assigning reads to their matching library member, all read counts were scaled to the median non-target read count for each time point. For library members with >30 initial read counts (t=0), read counts for each timepoint were normalized to the inital readcounts. Standard deviations in read counts were obtained by assuming Poisson-distributed read counts and propagating errors.

The experimental floor (defined as the median final timepoint value for all on-target controls) was first subtracted from all normalized read counts. Next, these adjusted normalized read counts were fit to a single exponential function to determine a cleavage rate (*k*_*clv*_): f(t) = e^(-*kclv* *t)^. Error in the fit was estimated by 50 rounds of bootstrapping with replacement.

### Single targets construction

Target sequences (Azenta, Table 3 primers Pr2.21, Pr6.77, Pr6.79, Pr7.09, Pr7.32) were amplified with Phusion Plus polymerase (Table 2 primers Pr6.29 and Pr6.30). Vector (Table 2 Pl0.63) was the same for each target and was PCR amplified (Phusion Plus polymerase, Table 2 primers Pr6.31 and Pr6.32). All reactions were column-purified (GeneJet column purification kit). Targets Δ3 and 3A4 had Cy3 and Cy5 labels added via PCR (Phusion Plus polymerase, Table 3 primers Pr7.50 and Pr8.27). Δ3 had both Cy3 and Cy5 simultaneously, while 3A4 had either one added at a time. Cy3 was always placed on a target strand and Cy5 on a non-target strand. All reactions were column-purified (GeneJet column purification kit).

**Table 3.**
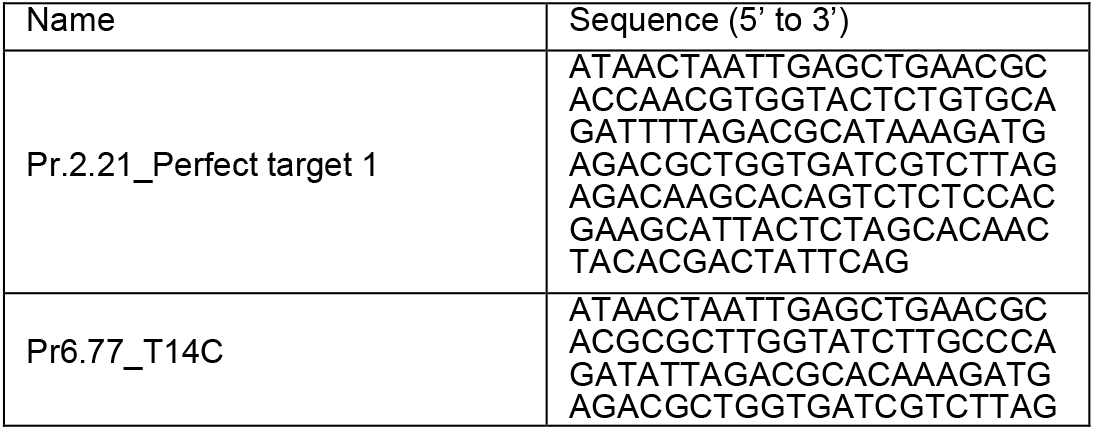
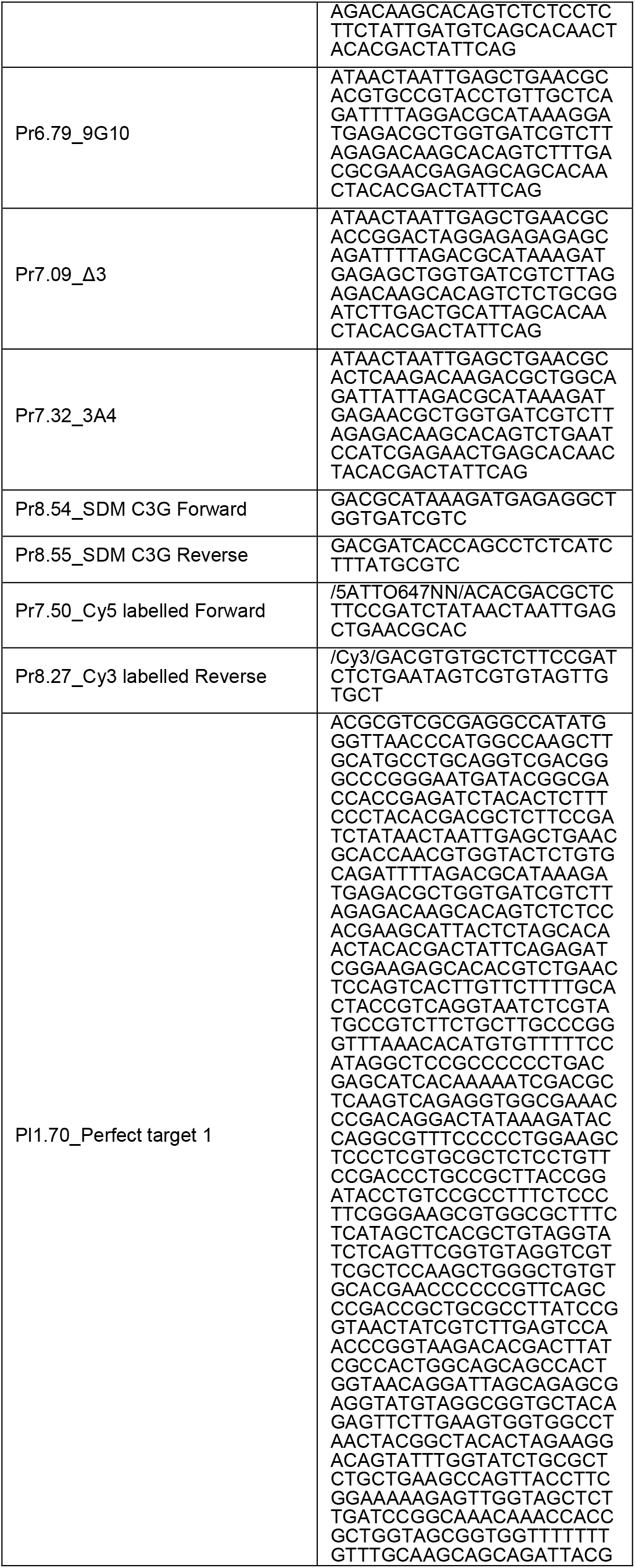
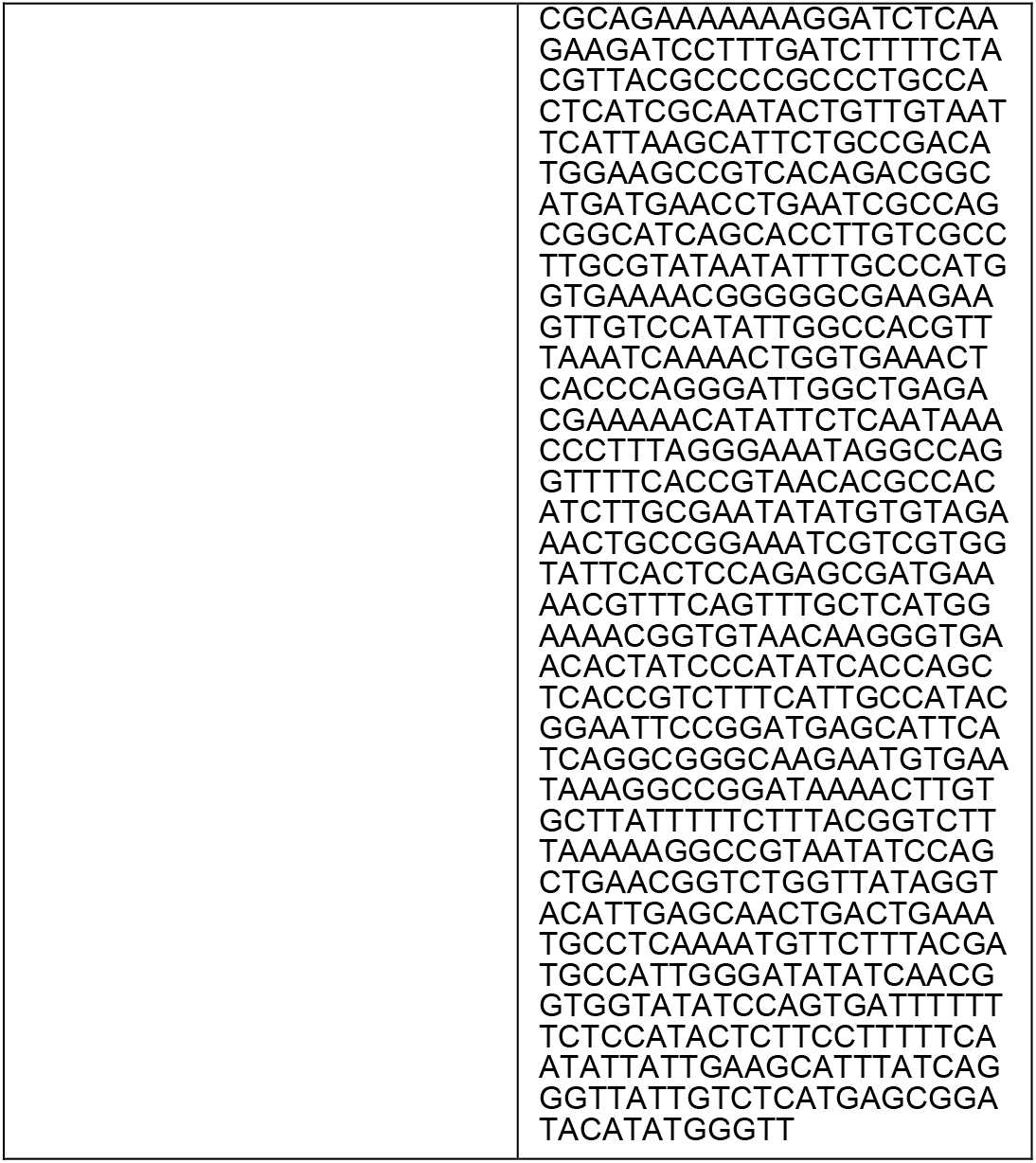
Single targets construction primers.

Plasmids were assembled using HiFi assembly (NEB, 5:1 insert to vector ratio) and electroporated (TOP10 competent cells, homemade). The plasmids were extracted from several randomly picked colonies (GeneJet plasmid miniprep kit, Thermo Scientific; Monarch plasmid miniprep kit, NEB) and sent out for sequencing (SeqVision UAB). Colonies with the correct assemblies were amplified (ZymoPURE II plasmid midiprep kit, Zymo Research).

Target C3G was made from ‘perfect’ on-target plasmid (Table 3 Pl1.70) by site-directed mutagenesis (Phusion Plus polymerase, 56°C annealing, Table 3 primers Pr8.54 and Pr8.55). The resulting reaction was treated with DpnI (Thermo Scientific) and electroporated (TOP10 competent cells, homemade). The plasmids were extracted from several randomly picked colonies (GeneJet plasmid miniprep kit, Thermo Scientific; Monarch plasmid miniprep kit, NEB) and sent out for sequencing (SeqVision UAB). Colonies with the correct assemblies were amplified (ZymoPURE II plasmid midiprep kit, Zymo Research). Half of each plasmid sample was relaxed with Topoisomerase I (NEB) or Nb. BsrDI (NEB). The relaxation was confirmed on a 1% Agarose gel electrophoresis in TBE buffer and the plasmids were then ethanol purified.

### Single target nicking and cleavage assays and analysis

The assays were performed on three types of DNA topologies for each of the selected single targets – supercoiled plasmid, relaxed plasmid, and linear PCR product. The assay was performed with Cas9 (NEB), gRNA (Table 1, R11) and DNA diluted in 100 mM NaCl, 50 mM Tris-HCl, 10 mM MgCl2, 100 µg mL−1 recombinant albumin, pH=7.9 (r3.1 buffer, NEB). After the RNP was formed by incubating Cas9 with an excess of gRNA for 15 minutes at room temperature, DNA was exposed to it for different amounts of time (0, 0.2, 0.5, 1, 3, 10, 30, 100, 300, ≥1000) before the reaction was stopped (60mM EDTA, 1U Proteinase K), and incubated at 37°C for 30 minutes. Some additional replicates omitted two final timepoints. The cleavage products were then visualized on 1% agarose (plasmid cleavage and nicking), 10% native PAGE (linear DNA cleavage) or 12% Urea PAGE in TBE buffer (linear DNA nicking).

To obtain cleavage rates from the gels, each gel was analyzed with GelAnalyzer 19.1 to quantify DNA band intensity. For fluorescently labelled targets, we quantified bands (∼200-180 bp) and cut products (∼90-68bp) after subtracting bands (∼90-85bp) present in the control. The parameters used on the program included background removal by a rolling ball with a peak width tolerance in 20% of lane profile length. Then, the values were normalized per lane and timepoint, and scaled to t=0. Finally, we obtained cleavage rate *k*_*clv*_ by fitting a single exponential function to the diminishing population of intact or nicked DNA: g(t) = e^(-*kclv* *t)^. We similarly obtained nicking rate *k*_*nck*_ by fitting a function to the diminishing population of intact DNA only: h(t) = e ^(-*knck* *t)^.

### Statistics

After obtaining cleavage rates, each set of single target assay replicates were tested for normal distribution using Shapiro-Wilk test, and the significance between tested conditions were analyzed using One-Way ANOVA and Tukey’s multiple comparison post-hoc test with Past v4.03^77^. If no cleaved products appeared in the last two timepoints, no cleavage rate was assigned to that target.

### Model methods

We fitted the mechanistic model CRISPRzip to the cleavage kinetics in library 1, as follows: First, we established a parameter set for this library on linear DNA, following the procedure described in Offerhaus et al. (2025)^42^. The landscape parameters (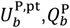 and *k*_f_) were trained to the relaxed-DNA cleavage and binding kinetics for all 1770 targets with up to 2 mismatches in 200 independent training runs. From the 50% best-scoring results, we calculated the median protein contributions (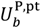and 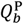), and determined the corresponding R-loop extension rate (*k*_f_) by optimization (Supp. Fig. 2).

Next, we estimated the plasmid torque level τ_0_ on the basis of the CRISPRzip torque parameters *θ* = 34° and *φ* = 50° estimated from torque spectroscopy.^34,42^ This torque-aware CRISPRzip version was trained to all supercoiled library-1 targets with ≥50 initial counts (1/1 perfect target, 60/60 single-mismatch targets, 851/1709 double-mismatch targets). Our modeling explicitly considered the coupling between the total R-loop opening angle *ω* and the plasmid torque τ,

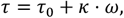

where τ_0_ is the initial plasmid torque (τ_0_ < 0 for negative supercoils), and *k* is the plasmid’s torsional stiffness. For our 1.7-kb plasmids (length *L* = 578 nm) with effective torsional persistence length *C*_eff_ = 20 nm,^78^ we used *k* = *C*_eff_⁄*L k*_B_*T* = 0.035 *k*_*B*_*T* rot^−1^ = 0.9 pNnm rad^−1^ . We performed a weighted least-squares fit of the plasmid torque ττ_0_ to the kinetic cleavage data, with cost function

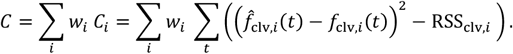

Here, *C*_i_ is the partial fit cost to target *i*, which scores the fit between the predicted cleaved fraction 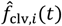 and the measured cleaved fraction *f*_clv,*i*_ (*t*). The value *C*_*i*_ compares this score to RSS_clv,*i*_, the residual sum of squares of individual exponential fit curves *y*_clv,*i*_ = 1 − exp−*k*_clv,*i*_ *⋅ t* to each target *i*. The target weights *w*_*i*_ were defined such that the perfect target (*w*_*i*_ = 1), single-mismatches (*w*_*i*_ = 1/60) and double-mismatches (*wi* = 1/851) all contributed equally to the fit cost. After obtaining a fit value for τ_0_, we expressed the corresponding cleavage predictions in terms of the effective cleavage rates *k*_clv, *i*_ obtained with an exponential fit to the predicted cleavage curves.

## Acknowledgements

We thank members of the Jones and Depken labs for valuable discussion, insights and editing of this manuscript. We thank Dr. Patrick Pausch and Dr. Ralf Seidel for insights and critical analysis of the manuscript. We thank the EMBL sequencing facility (GeneCore, Heidelberg, DE) for sequencing assistance.

H.S.O. acknowledges financial support from the Biononscience Department at TU Delft. This work was partially funded by the European Research Council (Project 101078247—PROTEGE; S.K.J.), the Ministry of Education, Science and Sports of the Republic of Lithuania (Program “University Excellence Initiatives” 12-001-01-01-01, project No. S-A-UEI-23-10; S.K.J.), and a European Molecular Biology Organization Installation Grant (Project IG 5728-2024; S.K.J.).

## Authorship statement

Conceptualization: IJ, HO, MD, SJ. Methodology: IJ, HO, MD, SJ. Software: HO, MV, MD, SJ. Validation: IJ, HO, UB. Formal analysis: IJ, HO, MV, UB. Investigation: IJ, HO, MV, UB. Resources: SJ, MD. Data curation: IJ, HO. Writing: IJ, HO, MD, SJ. Editing: IJ, HO, MV, MD, SJ. Supervision: IJ, SJ, MD. Project administration: MD, SJ. Funding acquisition: SJ, MD.

## Data and software availability

Raw sequencing data is available through European Nucleotide Archive database with the project accession number PRJEB90382. The NucleaSeq^15^ pipeline for processing NGS reads from cleavage experiments is available on Github (https://github.com/JonesLabEU/nucleaseq).

