## Supplementary material for "Supercoiling twists Cas9 off-target discrimination when nicking and cleaving": Supplmental Figures

### Supplementary figures

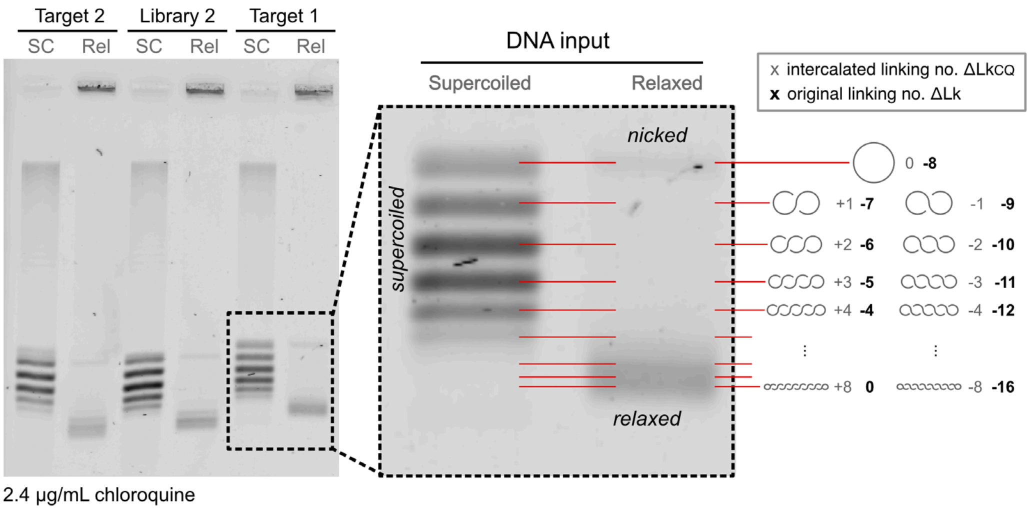


***Supplementary figure 1. Gel-based superhelical density determination.***

Illustration of supercoiling quantification method for Target 1 plasmids. The supercoiled and relaxed plasmid DNA inputs were ran on an agarose gel mixed with 2.4µg/mL of chloroquine in TBE buffer with the same concentration of chloroquine (for more detailed methods please refer to the methods section). Band positions in the gel arise from the linking number differences at the experimental chloroquine level, $\Delta\mathrm{Lk_{CQ}} = \Delta\mathrm{Lk} + f([\mathrm{CQ}])$. Here, the original linking number difference $\Delta\mathrm{Lk} = \mathrm{Lk} - L/h_0$ applies to a plasmid of length $L = 1.7$ kbp and B-DNA pitch $h_0 = 10.5$ bp in absence of chloroquine. The function $f([\mathrm{CQ}])$ captures how chloroquine increases DNA pitch beyond $h_0$. In the lane with the originally relaxed plasmids, the center of the intact-plasmid bands (labeled 'relaxed DNA input') revealed $\Delta\mathrm{Lk} = 0$, and the nicked plasmids in the population (labeled 'nicked') showed $\Delta\mathrm{Lk_{CQ}} = 0$. Based on the total band count, we estimated $f([\mathrm{CQ}]) = 8$. With this value, each band in the supercoiled (SC) population corresponded to two possible $\Delta\mathrm{Lk}$ values, as 1D electrophoresis cannot discern positive and negative supercoiling[48]. In line with previously measured $\sigma = -0.032 \pm 0.005$ for 2-50 kb plasmids in *E. coli* [79], we chose the lower value for $\Delta\mathrm{Lk}$. Next, we obtained the band intensity profiles (see methods) and estimated the distribution mean and standard deviation by maximum likelihood estimation. This approach resulted in $\Delta\mathrm{Lk} = -5.29 \pm 1.15$ for Target 2, $\Delta\mathrm{Lk} = -4.77 \pm 1.14$ for Library 2, and $\Delta\mathrm{Lk} = -5.28 \pm 1.31$ for Target 1. A weighted average of these three results yielded a superhelical density of $\sigma = \Delta\mathrm{Lk} \,/\, (L/h_0) = -0.033 \pm 0.007$.

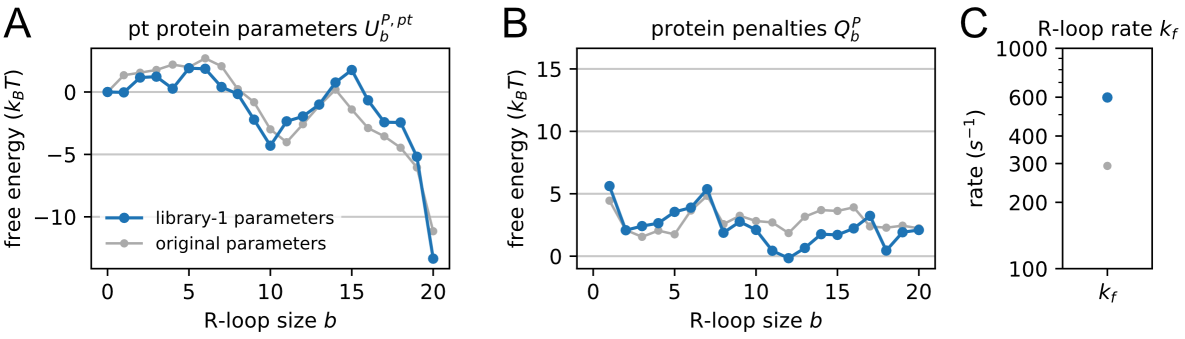


***Supplementary figure 2. Custom CRISPRzip parameter set.***

Library-1-specific CRISPRzip parameter values obtained for this study (blue, see Methods), alongside original parameter values (gray). The parameter sets comprise 20 perfect-target protein parameters $U_b^{\mathrm{P,pt}}$ (**A**), 20 protein penalties $Q_b^{\mathrm{P}}$(**B**) and 1 R-loop extension rate $k_\mathrm{f}$ (**C**).

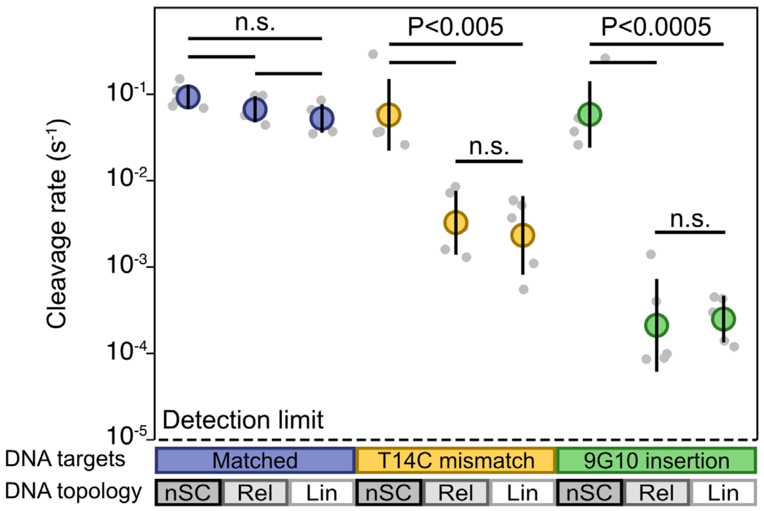


***Supplementary figure 3. Gel-obtained cleavage rates of supercoiled vs relaxed on- and off-targets.***

Data is present as mean±SD, N=5 in each group. One-Way ANOVA with Tukey's post-hoc test. nSC: negatively supercoiled, Rel: relaxed, Lin: linear.
